# Topographic, cognitive, and neurobiological profiling of the interdependent structural and functional connectome in the human brain

**DOI:** 10.1101/2024.01.21.576523

**Authors:** Xiaoyue Wang, Lianglong Sun, Xinyuan Liang, Tengda Zhao, Mingrui Xia, Xuhong Liao, Yong He

**Author notes:** These authors contributed equally (Xiaoyue Wang, Lianglong Sun).

## Abstract

The structural connectome (SC) is tightly coupled to the functional connectome (FC) in the human brain. Most previous related studies have modeled and analyzed SC or FC as isolated brain networks. However, challenges remain in modeling the interdependent structural-functional connectome and elucidating its cognitive implications and molecular underpinnings. Here, we present a multilayer connectome model composed of SC and FC components and further characterize their interacting topological properties. We found that the interdependent connectome is topographically heterogeneous, with the transmodal cortex exhibiting greater modular variability across layers. This spatial topography reflects cortical hierarchy and evolution and shows high test-retest reliability, reproducibility, and heritability. The interdependent connectome contributes to high-order cognitive processes and is associated with multiple neurotransmitter systems and transcriptional signatures of synaptic transmission. Our results provide insights into the nontrivial interdependencies of SC and FC, highlighting their cognitive significance and the molecular mechanisms underlying the connectome of connectomes.

## Introduction

The structural connectome (SC) and functional connectome (FC) are two indispensable components of the human brain connectome. The SC describes structural connectivity maps representing white matter pathways between regions ^1^, whereas the FC describes functional synchronization patterns representing temporal associations between regions ^2, 3^. These two connectomes interact and depend on each other to jointly maintain the functioning of the brain and further support cognitive processing. Elucidating the complex interplay between the SC and FC is one of the central challenges in network neuroscience. Previous empirical and computational modeling studies have consistently reported SC-FC coupling ^4-6^ or SC constraints on FC ^7-10^. Despite their importance, these previous studies have mainly modeled and analyzed SC or FC as isolated brain networks, ignoring the interdependent nature of the two connectomes.

Interdependent network theory ^11, 12^ provides an important mathematical framework for studying the connections between different types of networks. In recent years, research has been conducted to model and analyze the interdependent brain connectome that comprises SC and FC layers. For instance, several studies have employed integrated SC and FC features in the multilayer connectome to reveal nontrivial properties, such as overabundant network motifs or subgraphs ^13-16^, core hub regions ^17-19^, and core-periphery structures ^20, 21^. However, whether and how the SC and FC layers show different connectivity profiles in the interacting connectome are not yet well understood. To date, only one initial study has reported discordant global assortative mixing property between the SC and FC layers ^22^. However, how the SC and FC layers are topographically coordinated by different nodes in the interdependent connectome and how such multilayer coordination contributes to cognitive processes remain to be elucidated. Moreover, the neurobiological basis of the interdependent structural-functional connectome remains unknown. It is particularly important to answer these questions to better understand the organizational principles of interdependence in the unified structural-functional connectome and to elucidate the underlying biological mechanisms that govern the connectome.

A fundamental property of brain connectomes is that they exhibit a community or modular architecture that captures segregated and integrated processing ^23, 24^. Previous studies using isolated network models have shown that the SC and FC have different modular architectures, i.e., SC modules are anatomically constrained, whereas FC modules are functionally distributed ^5, 25, 26^. Currently, the multilayer modular organization of the interactive structural-functional connectome, particularly the brain nodes that are responsible for network communication between layered SC and FC modules, has not been well characterized. Connectome mapping studies have consistently shown a major primary-to-transmodal axis in the human brain ^27, 28^, with association cortices, including the default mode and frontoparietal regions, primarily involved in abstract cognition ^29^. Therefore, we reason that the transmodal cortices in the interdependent connectome play a major coordinated role between the layered SC and FC modules to promote cognitive diversity.

Previous SC or FC studies suggested that the spatial topography of the primary-to-transmodal axis aligns with that of neurobiological properties, such as gene expression ^30^, neurotransmitter receptor and transporter density ^31^. Studies have shown that gene expression related to synaptic functionality, including ion channel activity and synaptic functions (such as neurotransmitter release), is recapitulated within functional networks^32^. The late-developing heteromodal cortex manifests less genetic association than the primary cortex ^33, 34^. Neurotransmitters influence the balance between the integration and segregation of brain systems ^35^, and receptor distributions reflect structural and functional organization ^36^. Taken together, these previous studies based on isolated SC or FC networks have suggested potential genetic and molecular underpinnings of the modular architecture of the brain. Therefore, our second hypothesis is that the coordinated patterns of transmodal cortices across layered SC and FC modules are closely related to neurobiological properties, such as neurotransmitter receptors and transporters and gene expression profiles.

To address these issues, we investigated the interdependent structural-functional connectome of the human brain using resting-state functional magnetic resonance imaging (rs-fMRI) and diffusion MRI (dMRI) data from the Human Connectome Project (HCP) S1200 dataset ^37^. We first created a multiplex connectome comprising SC and FC layers and further identified the modular architectures using a multilayer modularity algorithm ^38, 39^. Next, we quantified the variability in the module affiliation of brain nodes between the SC and FC layers using multilayer modular variability ^40^. We then investigated whether the spatial topography of multilayer modular variability reflects the cortical gradient spanning from the primary region to the transmodal regions and further validated its reliability (using the HCP test-retest dataset), reproducibility (using the HCP half-split dataset), heritability (using the HCP twin-based dataset), and cognitive relevance (using the HCP cognitive dataset). Finally, to elucidate the neurobiological underpinnings of the coupled SC and FC connectomes, we conducted multivariate analysis to establish associations with neurotransmitter systems ^36^ and gene expression signatures ^41^, respectively.

## Results

### Construction of an interdependent structural-functional connectome using a multilayer network model

To model the interdependent brain connectome, for each individual, we reconstructed SC and FC connectomes using multimodal neuroimaging data from 1,012 healthy participants from the HCP S1200 dataset ^37^. Specifically, we defined network nodes based on a surface-based multimodal parcellation atlas ^42^ with 180 cortical areas per hemisphere (Fig. 1a). For network edges, we considered the Pearson correlation coefficient between the time series of all pairs of nodes for the FC ^26^ and the probabilistic diffusion tractography between nodes for the SC ^43^ (detailed in the Methods section). Considering that different ranges and properties of SC and FC connections could lead to disproportionate contributions to the following multilayer network analysis, we normalized the FC and SC matrices to a uniform range of 0-1 (Fig. 1b and 1c). We then modeled the interplay between the SC and FC in a multiplex framework that establishes interlayer connections based on direct correspondence between identical nodes. This process resulted in a two-layer interdependent SC-FC network for each individual (Fig. 1d), represented by a supra-adjacency matrix ^44^ where the diagonal blocks represent the intralayer connections and the off-diagonal blocks correspond to the interlayer connections. The multilayer modularity detection algorithm ^38, 39^ was applied to the interdependent structural-functional connectome, yielding a modular architecture that characterizes intra- and interlayer interactions simultaneously (Fig. 1e). To quantify how much the multilayer modular architecture in the interdependent structural-functional connectome differs between the SC and FC layers, we computed the multilayer modular variability of brain nodes based on their module assignments across layers ^40^. The greater the multilayer modular variability (e.g., node A in Fig. 1e), the greater the differences in the module structures to which nodes belong in the SC and FC layers.

**Fig. 1.**
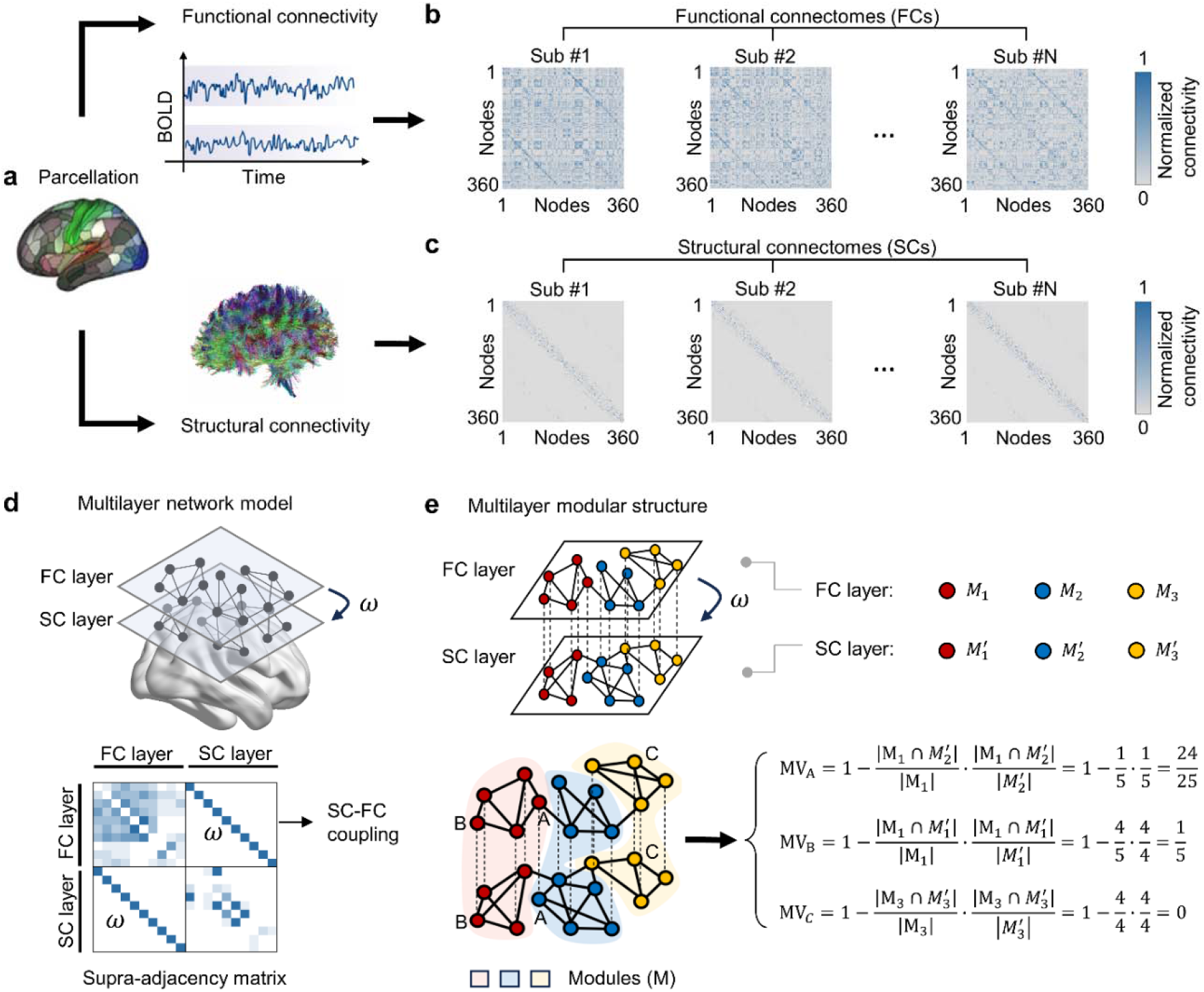
Framework for the construction and analysis of the interdependent SC-FC connectome. **a**, Multimodal parcellation (MMP1.0) was applied to define cortical areas, which were parcellated into 180 areas per hemisphere ^42^. **b, c** The FC and SC connectomes for each individual, wherein connectivity between each pair of cortical regions was obtained using Pearson correlation and probabilistic diffusion tractography, respectively. **d**, A schematic representation of the interdependent structural-functional connectome (top panel). The FC and SC form different layers, and dependency links between FC and SC were established by the multiplex coupling parameter w (w = 1), in which the corresponding nodes located in different layers were coupled in a one-to-one manner. The multilayer network can be represented by a supra-adjacency matrix ^44^, where the diagonal blocks represent the intralayer connections and the off-diagonal blocks correspond to the interlayer connections (bottom panel). **e**, The multilayer modularity detection algorithm ^38, 39^ was used to extract the modular architecture of the interdependent structural-functional connectome. Here, we exemplified the multilayer connectome as divided into three modules, with different modules rendered in different colors. The cross-layer module affiliation variability of nodes was evaluated using the multilayer modular variability (MV) metric ^40^. The calculation process of this metric is illustrated in detail in the figure, taking nodes A, B, and C as examples.

### The spatial topography of multilayer modular variability in the interdependent structural-functional connectome reflects cortical hierarchy and evolution

For each individual, we identified multilayer connectome modules and computed multilayer modular variability in the interdependent structural-functional connectome (Fig. 2a and 2b). The group-level multilayer modular variability showed substantial spatial heterogeneity across the cortex, with greater variability predominantly in the lateral prefrontal and parietal regions, dorsal medial prefrontal cortex and lateral temporal regions and less variability in the sensorimotor, visual, and ventral medial prefrontal cortex (Fig. 2c, left panel). Furthermore, we investigated whether the spatial pattern of multilayer modular variability represents cortical hierarchical organization. First, we stratified the 360 cortical regions into four hierarchies illustrating a transition from primary sensory regions to the transmodal cortex ^45^. The heteromodal system (spin test *P* value (P_spin_) < 0.001) exhibited significantly greater multilayer modular variability than did the null models, while the primary (P_spin_ = 0.0003) and unimodal (P_spin_ < 0.001) systems exhibited significantly less multilayer modular variability (Fig. 2c, right panel). Second, we found that the topographic organization of multilayer modular variability significantly correlated with a well-established macroscale connectome gradient architecture from unimodal to transmodal ^28^ (Pearson’s r (358) = 0.56, P_spin_ < 0.0001, confidence interval (CI)□=□[0.48, 0.62], two-tailed; Fig. 2d). Given that greater multilayer modular variability was observed in the association regions that are thought to be phylogenetically late-evolving regions, we hypothesized that there would be a potential evolutionary root for the interaction of the structural-functional connectome. By correlating with the cortical evolutionary expansion ^46^, we found that highly conserved sensory areas exhibited relatively less SC-FC modular variability, while highly expanded transmodal areas exhibited greater SC-FC modular variability (Pearson’s r (178) = 0.51, P_spin_ < 0.001, CI = [0.39, 0.61], two-tailed; Fig. 2e). Taken together, these results suggested that the modular topography of the multilayer structural-functional connectome is regionally heterogeneous, reflecting primary-to-transmodal organization and a cortical evolutionary mechanism.

**Fig. 2.**
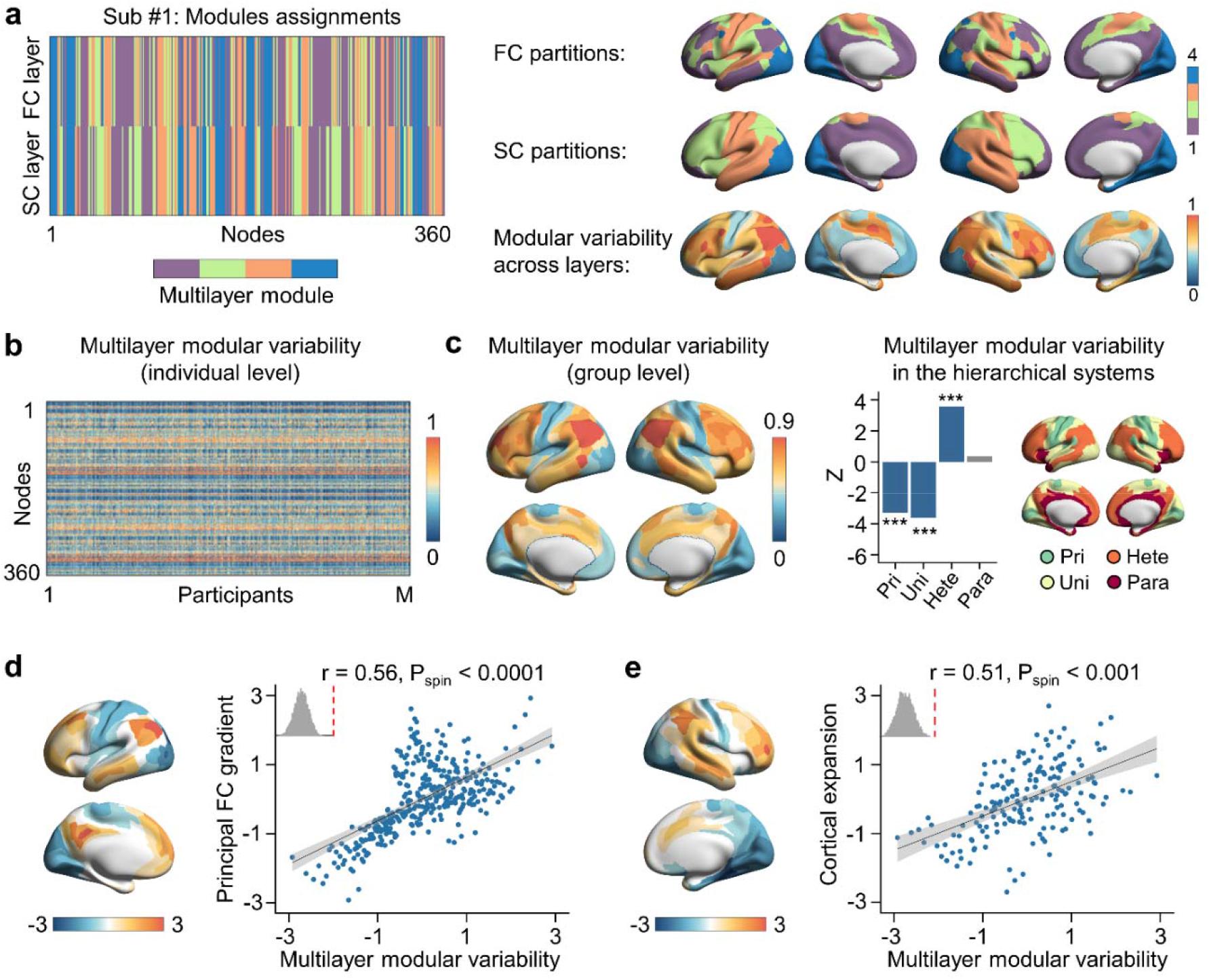
The spatial topography of multilayer modular variability in the coupled structural-functional connectome reflects the cortical gradient and evolutionary expansion. **a**, The multilayer modular structure of the interdependent structural-functional connectome for a representative participant, in which the elements of the matrix represent the modules to which the nodes were assigned in the FC and SC layers (left panel). The partitions of the FC and SC are projected onto the brain cortex, and based on this, the cross-layer module affiliation of each node is tracked to obtain the spatial distribution of the participant’s multilayer modular variability (right panel). **b**, Multilayer modular variability at the individual level. **c**, The spatial topography of group-level multilayer modular variability (left panel), in which the heteromodal system (P_spin_ < 0.001) exhibited significantly greater multilayer modular variability than did the null models, while the primary (P_spin_ = 0.0003) and unimodal (P_spin_ < 0.001) systems exhibited significantly less multilayer modular variability (right panel). Nodewise multilayer modular variability values were averaged according to their hierarchical systems. The mean multilayer modular variability of the system-specific is expressed as a z score relative to the null model (spin test 10,000 repetitions), in which positive (negative) z values indicate larger (less) multilayer modular variability than expected by chance. The statistically significant and nonsignificant systems are shown in color and gray, respectively. The multilayer modular variability was significantly (spin test 10,000 repetitions) associated with the functional connectivity gradient (Pearson’s r (358) = 0.56, P_spin_ < 0.0001, confidence interval (CI)□=□[0.48, 0.62], two-tailed) (**d**) and evolutionary expansion of cortical surface area (Pearson’s r (178) = 0.51, P_spin_ < 0.001, CI = [0.39, 0.61], two-tailed) (**e**). The gray shaded envelopes in the scatter plots indicate the 95% CI, the upper left corners of the scatter plots show the histograms of r values obtained from the null model, and the vertical red dotted lines denote the empirical r values. To better visualize the scatter plots, the raw values (including those for the cortical gradient, cortical expansion and multilayer modular variability) were scaled using a rank-based inverse Gaussian transformation ^47^. The brain maps were generated using the BrainNet Viewer package ^48^ on the inflated cortical 32K surface ^42^. Pri, primary cortex; Uni, unimodal cortex; Hete, heteromodal cortex; Para, paralimbic cortex. *** P < 0.001.

### Reliability, reproducibility, and heritability of multilayer modular variability in the interdependent structural-functional connectome

Having demonstrated that multilayer modular variability in the interdependent structural-functional connectome has a specific topographic distribution across the cortical hierarchy, we next examined whether this distribution is test-retest reliable, reproducible and heritable.

First, we used the HCP Test-Retest dataset to evaluate test-retest reliability. This dataset included 42 participants (aged 30.4 ± 3.33 years, 30 females) who underwent a second MRI scanning session scheduled between 0.5 and 11 months after their first session. Specifically, we calculated the Pearson correlation coefficient of multilayer modular variability between the same individuals in different sessions and between different individuals. We found that the intraindividual spatial similarity of multilayer modular variability (Pearson’s r: 0.69 ± 0.123) was significantly greater (nonparametric permutation test *P* value (P_perm_) < 0.0001) than the interindividual similarity (Pearson’s r: 0.48 ± 0.065; Fig. 3a). Furthermore, for each brain node, we performed the intraclass correlation (ICC) ^49^ analysis to estimate its test-retest reliability of the multilayer modular variability in the interdependent structural-functional connectome. The highest test-retest reliability was in the dorsolateral prefrontal and inferior parietal cortex (ICC > 0.6) (Fig. 3b, left panel). Compared to those of the other systems, the ICC of the heteromodal system was significantly greater than those of the null model (P_spin_ < 0.001), while the ICC of the paralimbic system was significantly lower than the level expected by chance (P_spin_ = 0.0039) (Fig. 3b, middle panel). The spatial distribution of the ICC map showed a significant correlation with that of the multilayer SC-FC modular variability map (Pearson’s r (358) = 0.23, P_spin_ < 0.0003, CI = [0.13, 0.32], two-tailed; Fig. 3b, right panel).

**Fig. 3.**
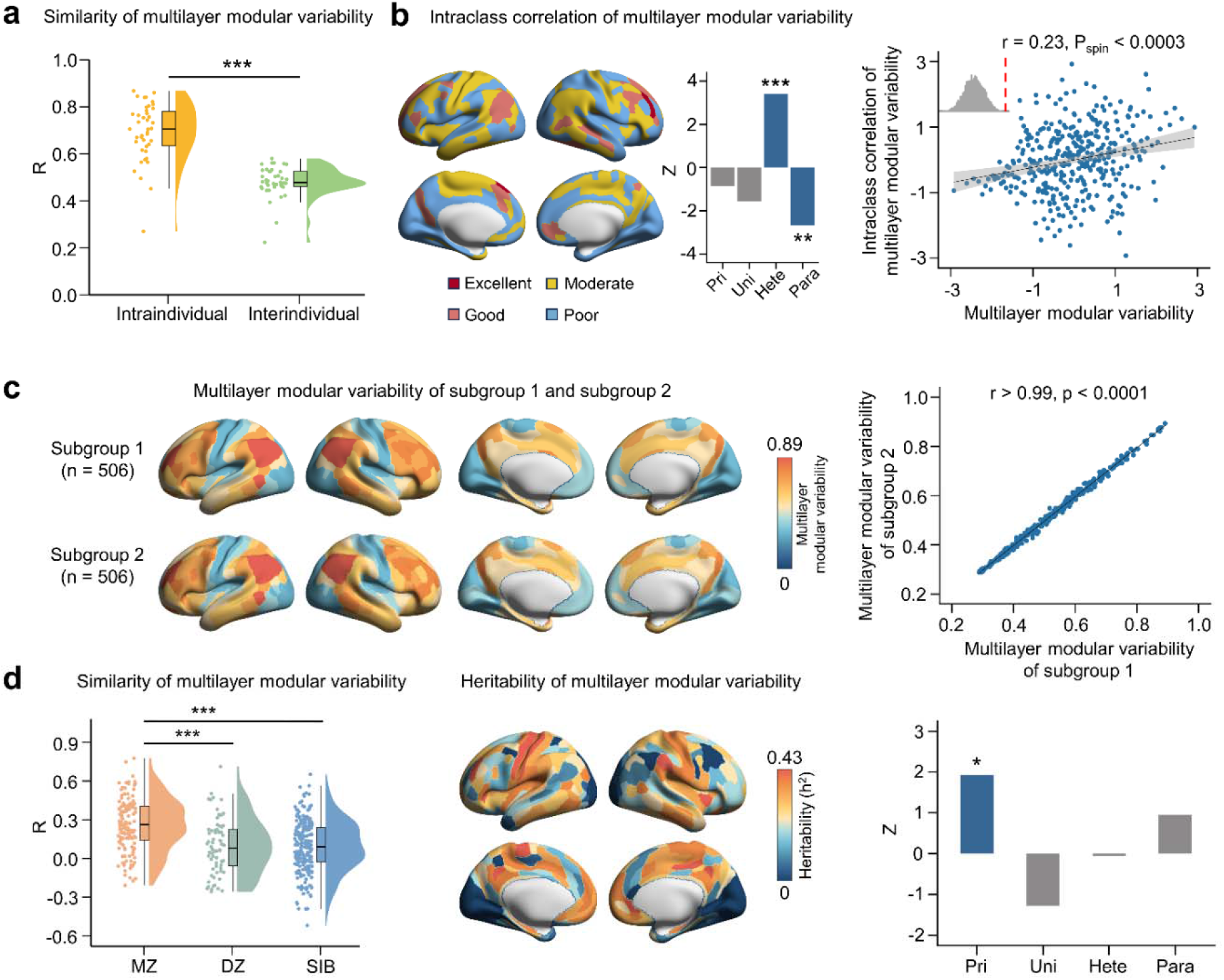
Multilayer modular variability is reliable, reproducible, and heritable. **a**, The intraindividual similarity of multilayer modular variability is greater than the interindividual similarity (P_perm_ < 0.0001, nonparametric permutation test 10,000 repetitions). **b**, The spatial pattern of the intraclass correlation (ICC) of multilayer modular variability (left panel), in which the heteromodal system exhibits a greater ICC (P_spin_ < 0.001) while the paralimbic system exhibits a lower ICC (P_spin_ = 0.0039) than null models (spin test 10,000 repetitions; middle panel), is shown. The ICC of multilayer modular variability was correlated with multilayer modular variability (Pearson’s r (358) = 0.23, P_spin_ < 0.0003, confidence interval (CI) = [0.13, 0.32], two-tailed, spin test 10,000 repetitions; right panel). The gray shaded envelope in the scatter plot indicates the 95% CI, the upper left corner of the scatter plot shows the histogram of r values obtained from the null model, and the vertical red dotted line denotes the empirical r value. To better visualize the scatter plot, the values of the raw variables were scaled using a rank-based inverse Gaussian transformation ^47^. **c**, One of the random splits of the 1000 iterations using a half-split strategy suggested that the group-level multilayer modular variability patterns were highly similar (Pearson’s r > 0.99, p < 0.0001). **d**, The similarity of multilayer modular variability between MZ pairs was greater (nonparametric permutation test, 10,000 repetitions) than that between DZ (P_perm_ < 0.0001) and sibling (P_perm_ < 0.0001) pairs (left panel). By estimating the regional heritability of multilayer modular variability (middle panel), the primary system was highly heritable (P_spin_ = 0.016, spin test 10,000 repetitions; right panel). The bounds of the boxplots in **a** and **d** represent the 1st (25%) and 3rd (75%) quartiles, the centerline represents the median, and the whiskers represent the minima and maxima of the distribution. The violin plots in **a** and **d** show the distribution of Pearson’s r values in the different groups indicated on the x axis. The bar plots in **b** and **d** show that the nodewise ICC (heritability) values were averaged according to their hierarchical systems. The mean ICC (heritability) of the system is expressed as a z score relative to the null model, in which positive (negative) z values indicate that the ICC (heritability) is greater (less) than expected by chance. Pri, primary cortex; Uni, unimodal cortex; Hete, heteromodal cortex; Para, paralimbic cortex; MZ, monozygotic; DZ, dizygotic; SIB, sibling. * P < 0.05, ** P < 0.01, *** P < 0.001.

Second, we validated the reproducibility of multilayer SC-FC modular variability. Specifically, we randomly (n = 1,000 repetitions) divided the 1,012 participants in the HCP S1200 dataset into two cohorts (subgroup 1 and subgroup 2). We found that the group-level multilayer modular variability was highly correlated in the interdependent structural-functional connectomes of the two subgroups (Pearson’s r = 0.994-0.999, p < 0.0001; Fig. 3c shows the results of one of the 1,000 repetitions), suggesting high reproducibility.

Finally, using twin and family data from the HCP S1200 dataset, which consists of 268 monozygotic (MZ) twins, 140 dizygotic (DZ) twins, 107 singletons, and 494 nontwins, we examined whether the interindividual similarity of multilayer modular variability differed between unrelated and genetically related individuals. We used Pearson correlation to measure the similarity of multilayer modular topography across participants. We found that the similarity varied for MZ twins (Pearson’s r: 0.26 ± 0.204), DZ twins (Pearson’s r: 0.10 ± 0.217), and siblings (Pearson’s r: 0.10 ± 0.196), with significantly greater similarity among MZ twins than among DZ twins (P_perm_ < 0.0001) and siblings (P_perm_ < 0.0001) (Fig. 3d, left panel). We further performed a twin-based heritability analysis to examine the heritability of the multilayer network modules in the interacting structural-functional connectome. We found that genetic factors exerted a regionally variable influence on the multilayer modular variability, with higher heritability observed in the somatosensory, lateral temporal, medial prefrontal, and parietal regions and lower heritability in the lateral frontal and parietal regions and visual cortices (Fig. 3d, middle panel). Similarly, we found that the heritability was not uniform across the four hierarchical systems, with the primary system being significantly more heritable relative to the null models (P_spin_ = 0.016, Fig. 3d, right panel). Taken together, our results indicated that the degree of spatial variability of multilayer module organization across the layered SC and FC is heritable.

### Multilayer modular variability in the interdependent SC-FC connectome is associated with high-order cognitive processes

We first examined whether the multilayer modular variability in the interdependent structural-functional connectome was spatially associated with neurocognitive flexibility quantified by the number of cognitive components proposed by Yeo et al ^50^. For each brain node, we calculated its neurocognitive flexibility by averaging the number of cognitive components of all voxels within that node (Fig. 4a). We found a significant correlation between multilayer modular variability and neurocognitive flexibility (Pearson’s r (358) = 0.27, P_spin_ = 0.004, CI = [0.17, 0.36], two-tailed; Fig. 4b). Based on the number of cognitive components involved, we categorized all brain nodes into four types: low flexibility (0 ≤ number of components < 1), moderate flexibility (1 ≤ number of components < 2), good flexibility (2 ≤ number of components < 3) and high flexibility (number of components ≥ 3). We found that the high-flexibility nodes exhibited high multilayer modular variability relative to the other types of nodes (Kruskal-Wallis test, Bonferroni correction, p < 0.001; Fig. 4c). These results suggested that brain nodes with higher multilayer modular variability tended to participate in multiple cognitive components and contributed to higher cognitive flexibility.

**Fig. 4.**
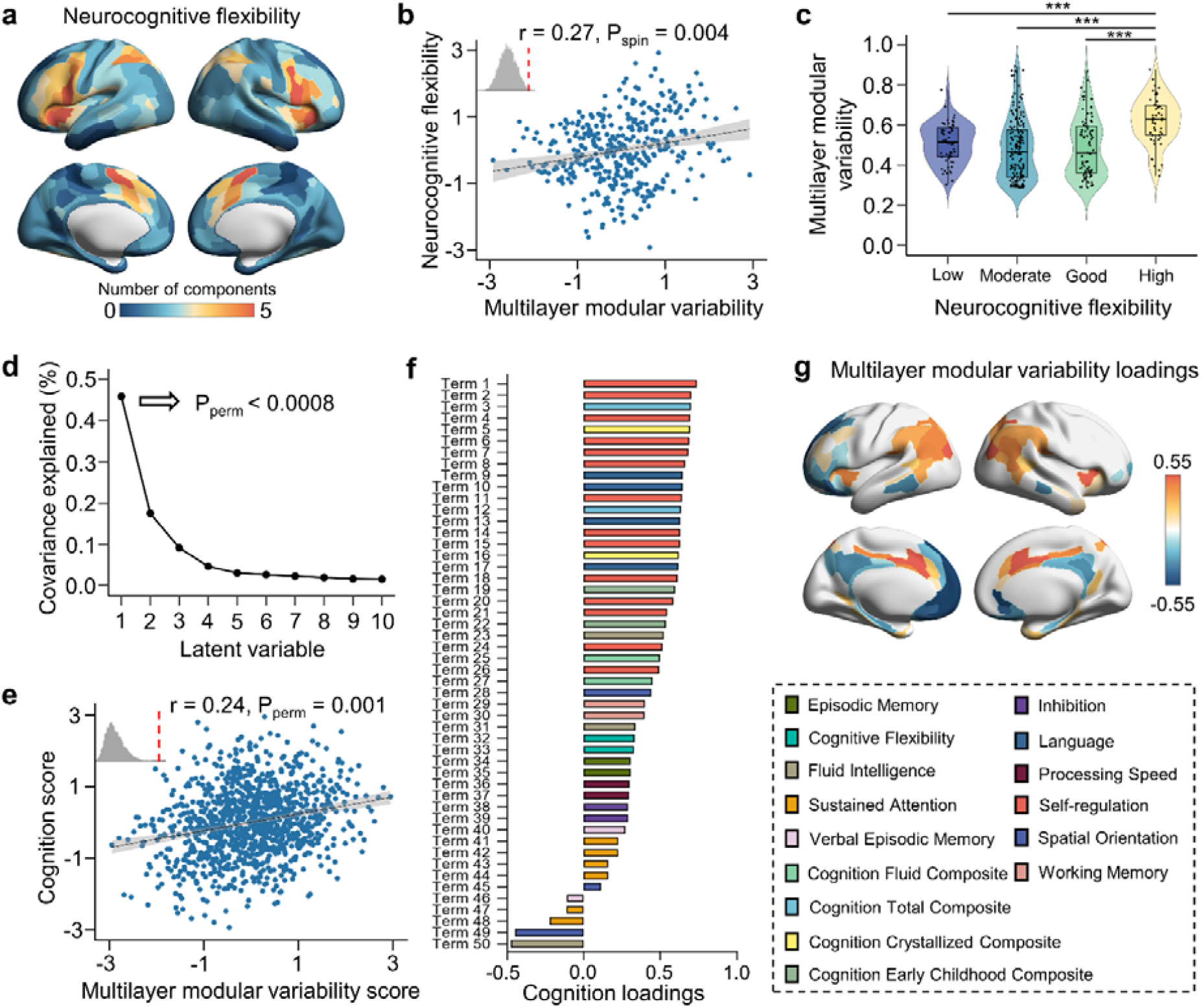
Associations between multilayer modular variability and high-order cognitive function. **a**, The neurocognitive flexibility of cortical regions is characterized by the number of cognitive components they engage in ^50^. **b**, Multilayer modular variability was significantly correlated with neurocognitive flexibility (Pearson’s r (358) = 0.27, P_spin_ = 0.004, confidence interval (CI) = [0.17, 0.36], two-tailed, spin test 10,000 repetitions). **c**, Nodes were categorized into four types, namely, low flexibility (0 ≤ number of components < 1), moderate flexibility (1 ≤ number of components < 2), good flexibility (2 ≤ number of components < 3), and high flexibility (number of components ≥ 3) nodes. The results of Kruskal-Wallis test indicated that nodes with high flexibility exhibited significantly greater multilayer modular variability (Bonferroni correction, p < 0.001). **d**, The first latent variable (LV1) can significantly account for 46% of the covariance between multilayer modular variability and cognition (P_perm_ < 0.0008, nonparametric permutation test 10,000 repetitions). **e**, For LV1, the multilayer modular variability score and cognition score were significantly correlated (Pearson’s r (1010) = 0.24, P_perm_ = 0.001, CI = [0.19, 0.30], two-tailed). **f** and **g** show the loadings of cognition terms and brain regions with significant contributions (bootstrapping resampling 1,000 repetitions). The cognitive processes depicted by these cognition terms are shown in the right panel with different colors. For detailed cognitive processes and cognitive loadings, see Supplementary Table 1. The gray shaded envelopes in the scatter plots indicate the 95% CI, the upper left corners of the scatter plots show the histograms of r values obtained from the null model, and the vertical red dotted lines denote the empirical r values. To better visualize the scatter plots, the raw variable values were scaled using a rank-based inverse Gaussian transformation ^47^. *** p < 0.001.

Next, we sought to investigate whether the multilayer modular variability in the interdependent structural-functional connectome is related to individual’s cognitive function. We applied multivariate partial least squares (PLS) analysis to separately estimate the extent to which the multilayer modular variability in the primary and transmodal cortices was related to cognitive performance. Specifically, we first stratified the cerebral cortex into the low-order area (consisting of primary and unimodal regions, 176 regions in total) and high-order transmodal area (consisting of heteromodal and paralimbic regions, 184 regions in total). PLS analysis revealed that there was no significant relationship between multilayer modular variability in the low-order cortex and cognitive performance. In contrast, for the transmodal cortex, the first latent variable (LV1) significantly (P_perm_ < 0.0008) captured 46% of the covariance between multilayer modular variability and cognition (Fig. 4d). Under the LV1, the multilayer modular variability score was significantly correlated with the cognition score (Pearson’s r (1010) = 0.24, P_perm_ = 0.001, CI = [0.19, 0.30], two-tailed; Fig. 4e). This correlation was determined by the brain regions and cognitive terms that contribute most to the latent variable. Therefore, we computed the loadings to determine the degree of contribution of each variable to the latent component and assessed the reliability of the brain region and cognitive term loadings through bootstrapping resampling (1,000 repetitions). For multilayer modular variability, regions with large positive loadings were located mainly in the inferior parietal cortex, temporal-parietal-occipital junction, and anterior cingulate cortex, whereas regions with large negative loadings were located mainly in the medial prefrontal, posterior cingulate, and lateral temporal cortices (Fig. 4g). Interestingly, we found that almost all the cognitive terms had positive loadings with terms belonging to self-regulation, cognition total composite, and cognition crystallized composite cognitive processes showing the largest loadings (Fig. 4f). These results demonstrated that greater multilayer modular variability in brain regions with positive loadings was associated with better high-level cognitive performance.

### The multilayer modular organization of the interdependent structural-functional connectome is predicted by neurotransmitter receptors and transporters

A previous study has shown that the spatial topography of neurotransmitter systems reflects the organizational architecture of brain networks ^36^. Here, we sought to investigate whether the multilayer modular organization in the interdependent SC-FC network is associated with neurotransmitter receptors and transporters. To do this, we obtained cortical distribution data of 19 neurotransmitter receptors and transporters from nine neurotransmitter systems provided by Hansen et al ^36^. Then, we calculated the average density of each cortical region in the Glasser360 atlas for each of the 19 receptors and transporters. We found that 4 out of the 19 receptor and transporter density distributions were significantly positively correlated with multilayer modular variability (Benjamini–Hochberg false discovery rate (FDR) correction, q < 0.05), namely, MOR (Pearson’s r (358) = 0.38, P_spin_ < 0.0001, CI = [0.28, 0.46], two-tailed), CB_1_ (Pearson’s r (358) = 0.29, P_spin_ < 0.0002, CI = [0.20, 0.38], two-tailed), 5-HT_4_ (Pearson’s r (358) = 0.20, P_spin_ = 0.0087, CI = [0.10, 0.30], two-tailed) and *α*_*4*_*β*_*2*_ (Pearson’s r (358) = 0.20, P_spin_ = 0.0042, CI = [0.10, 0.30], two-tailed) receptors (Fig. 5a and 5b). We further sought to explore the extent to which multilayer modular variability can be explained by receptor and transporter data. Using the multivariate elastic net regression model (*λ* = 0.011; Fig. 5c), we found that the spatial pattern of multilayer modular variability could be significantly predicted by the density distributions of neurotransmitter receptors and transporters (Pearson’s r (358) = 0.59, P_spin_ < 0.0001, CI = [0.52, 0.66], two-tailed; Fig. 5d). Moreover, 11 out of the 19 receptors and transporters significantly contributed to the prediction model (Fig. 5e); the highest contributions were from the MOR, 5-HT_4_, and *α*_*4*_*β*_*2*_ receptors. Together, our results highlighted the tight link between the interdependent structural-functional connectome and multiple neurotransmitter systems.

**Fig. 5.**
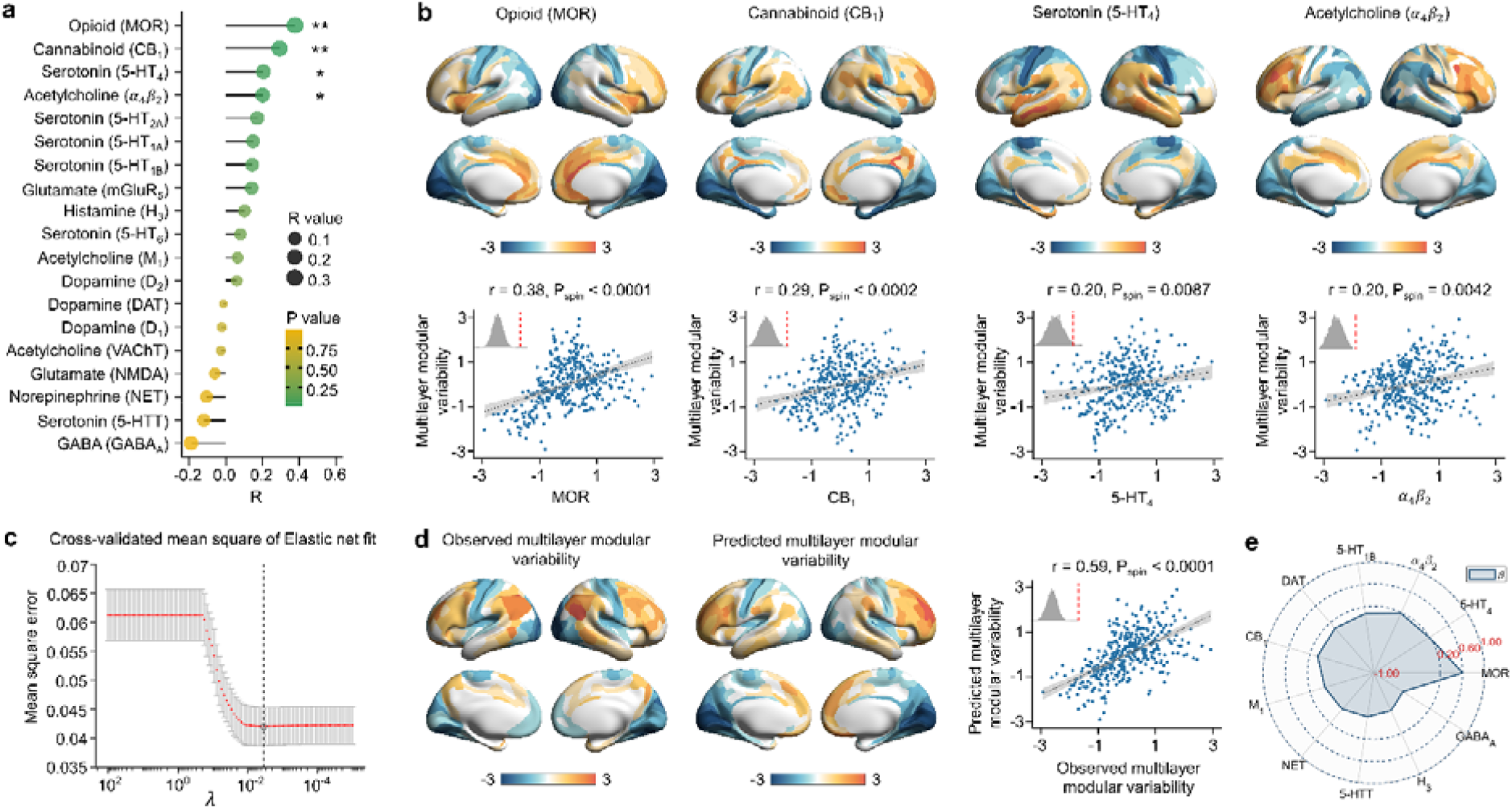
Associations between the spatial topography of multilayer modular variability and neurotransmitter receptor and transporter distributions. **a**, The density distributions of 4 out of 19 receptors and transporters were significantly correlated with multilayer modular variability after Benjamini–Hochberg false discovery rate (FDR) correction (P_spin_ < 0.05, spin test 10,000 repetitions). The spatial distributions and correlations of these receptors are shown in (**b**) (MOR: Pearson’s r (358) = 0.38, P_spin_ < 0.0001, confidence interval (CI) = [0.28, 0.46], two-tailed; CB_1_: Pearson’s r (358) = 0.29, P_spin_ < 0.0002, CI = [0.20, 0.38], two-tailed; 5-HT_4_: Pearson’s r (358) = 0.20, P_spin_ = 0.0087, CI = [0.10, 0.30], two-tailed; *α*_*4*_*β*_*2*_: Pearson’s r (358) = 0.20, P_spin_ = 0.0042, CI = [0.10, 0.30], two-tailed). **c**, The 10-fold cross-validated elastic net regression was performed with different *λ* values (100 values from 10^−5^ to 10^2^). The vertical black dotted line denotes the optimal *λ* values (*λ* = 0.011) with the minimum mean square error (MSE = 0.043). **d**, The observed and predicted multilayer modular variabilities are significantly correlated (Pearson’s r (358) = 0.59, P_spin_ < 0.0001, CI = [0.52, 0.66], two-tailed, spin test 10,000 repetitions). The gray shaded envelopes in the scatter plots indicate the 95% CI, the upper left corners of the scatter plots show the histogram of r values obtained from the null model, and the vertical red dotted lines denote the empirical r values. Elastic net regression provided a sparse output, in which 11 receptors and transporters significantly contributed to the prediction model, and the regression coefficient (*β*) of each receptor/transporter is shown in (**e**). For better visualization, the raw values (including receptor density and multilayer modular variability) were scaled using a rank-based inverse Gaussian transformation ^47^.

### Transcriptomic profiles are associated with multilayer modular architecture in the interdependent structural-functional connectome

Gene expression regulates the coordinated activity of neuronal populations and further shapes complex cognitive processes ^51^. Using regional microarray expression data from the Allen Human Brain Atlas (AHBA) dataset (n = 6, donor brains) ^41^, we investigated whether the multilayer module configuration in the interdependent structural-functional connectome was associated with gene expression profiles. PLS regression analysis revealed that the first PLS (PLS1) component, which explained 21.25% of the variance in multilayer SC-FC modular variability (P_spin_ = 0.02; Fig. 6a), exhibited a significant positive correlation between multilayer modular variability and regional gene expression (Pearson’s r (130) = 0.46, P_spin_ = 0.02, CI = [0.31, 0.59], two-tailed; Fig. 6b). The PLS1 component represented a gene expression profile with high expression mainly in the lateral frontal and parietal cortices but low expression in the sensorimotor and visual cortices. We then performed Gene Ontology (GO) enrichment analysis on genes associated with the transcriptome features of the PLS1 component. We found that genes ranked by weight from most positive to most negative were enriched in biological processes related to chemical synaptic transmission (FDR-corrected, q < 0.05; Fig. 6c, middle panel; Supplementary Table 2) and cellular components related to synapse part, plasma membrane, neuron part, transport vesicle, and secretory vesicle (FDR-corrected, all q < 0.05; Fig. 6c, right panel; Supplementary Table 2). No significant enrichment was observed for molecular function. We also performed GO enrichment analysis on the inverse ranking of genes. The significant enrichment terms are shown in Supplementary Table 3. Collectively, these results revealed a potential molecular basis for the multilayer module organization in the interacting structural and functional connectome.

**Fig. 6.**
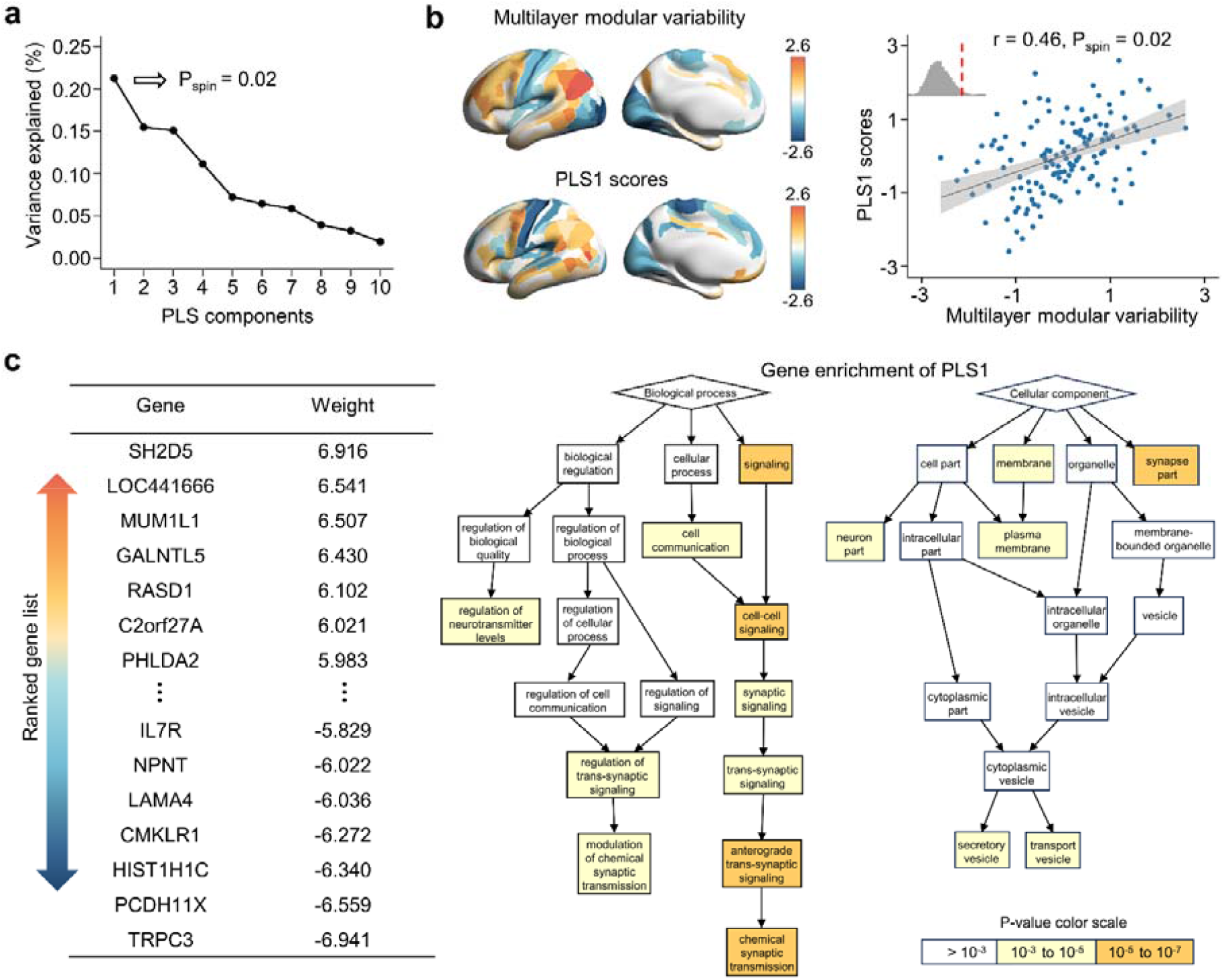
Association between the spatial topography of multilayer modular variability and gene expression profiles. **a**, PLS regression analysis results suggested that the first principal component (PLS1) of the gene expression matrix significantly captured 21.25% of the variance in multilayer modular variability (P_spin_ = 0.02, spin test 10,000 repetitions). **b**, The PLS1 scores and multilayer modular variability were significantly correlated (Pearson’s r (130) = 0.46, P_spin_ = 0.02, confidence interval (CI) = [0.31, 0.59], two-tailed, spin test 10,000 repetitions). The gray shaded envelope in the scatter plot indicates the 95% CI, the upper left corner of the scatter plot shows the histogram of r values obtained from the null model, and the vertical red dotted line denotes the empirical r value. To better visualize the scatter plot, the raw variable values were scaled using a rank-based inverse Gaussian transformation ^47^. **c**, The listed genes were ranked in descending order according to weight, which represents the contribution of each gene to the PLS1 component (left panel). Gene Ontology enrichment analysis of this gene list revealed that the genes were significantly (FDR-corrected, all q < 0.05) enriched in biological processes related to chemical synaptic transmission (middle panel) and cellular components related to synapse part, the plasma membrane, neuron part, transport vesicle and secretory vesicle (right panel).

### Sensitivity and robustness analysis

Head motion has long been thought to have profound effects on brain imaging data ^52, 53^. To validate the effect of head motion, we excluded participants with high head motion and then repeated our main analyses. Briefly, we measured each participant’s head motion indices by calculating the mean and mean absolute deviation of the frame-to-frame displacements from both the fMRI and dMRI scanning sessions and then excluded participants whose head motion indices exceeded 1.5 times the interquartile range of the corresponding index distribution ^10^.

Consequently, we completely excluded 95 out of 1,012 participants in the HCP S1200 dataset and 5 out of 42 participants in the HCP Test-Retest dataset. We then performed validation analysis with the remaining participants. We observed that the multilayer modular variability pattern in the interdependent structural-functional connectome after removing the participants with high-level motion was highly similar to our main results (Pearson’s r > 0.99, p < 0.0001; Supplementary Fig. 1). We then validated the reliability, reproducibility, and heritability of multilayer modular variability and found consistent results (Supplementary Fig. 2). The results of association analyses with cognitive data (Supplementary Fig. 3), neurotransmitter receptor and transporter data (Supplementary Fig. 4), and gene expression profiles (Supplementary Fig. 5) were also highly consistent with our main findings. All these validations suggested that our main findings are robust and are not affected by head motion.

Next, to evaluate the threshold effects on the multilayer modular properties in the interdependent structural-functional connectome, we applied different connection thresholds to the FC matrices. We observed that the spatial patterns of multilayer modular variability were highly reliable across different thresholds (both Pearson’s rs > 0.99, p < 0.0001) (Supplementary Fig. 6 and 7).

## Discussion

Using multimodal neuroimaging data and a multilayer network model, we constructed the interdependent structural-functional connectome and identified a multilayer modular architecture. We showed that the spatial topography of module variability across the SC and FC layers follows a primary-to-transmodal axis and that this pattern is test-retest reliable, reproducible, and heritable. We further showed that greater multilayer modular variability in the transmodal cortex contributes to greater cognitive diversity and abstract cognitive processes. Finally, the multilayer modular variability in the interdependent structural-functional connectome is closely associated with neurotransmitter receptor and transporter density and gene expression profiles, suggesting the neurobiological underpinnings of connectome.

Previous studies have reported heterogeneous correspondence between the SC and FC across the cortex, with high correspondence in the primary cortex and low correspondence in the association cortex ^4-8, 54-57^. Our results extended these findings by highlighting the interdependencies between the SC and FC in a multiplex framework. In the interdependent structural-functional connectome, the spatial variability pattern of module organization across layers aligns with a primary-to-transmodal axis. Previous studies have demonstrated that various brain properties, such as functional connectivity ^28^, gene expression ^30^, cognition ^58^, and receptors ^31^, follow this core organizational axis of the cerebral cortex ^27^. We have shown that the transmodal cortex has a markedly divergent correspondence between the layered SC and FC modules, which may be due to the fact that these areas are not bound by structural constraints.

Rapid expansion of the cortical mantle leads to local microcircuitry reorganization, effectively freeing the associative cortex from the strong constraints imposed by early activity cascades ^6, 34^. Thus, weaker structural constraints in the transmodal cortex allow for a more flexible modular architecture ^59, 60^. Notably heterogeneous communication preferences across the cortex, with unimodal regions communicating primarily at local scales and multimodal regions communicating primarily at global scales ^61^, provide another possible explanation for the observed divergence of the SC and FC. The primary sensory cortex is involved mainly in unitary neural circuitry and supports simple sensory functions, whereas the transmodal cortex mainly receives and integrates information from multiple sensory modalities and other heteromodal regions, resulting in more spatially distributed connection patterns to support global communication and integration between functional systems. Thus, the high variability of the multilayered module organization in the transmodal cortex supports the involvement of more extensive modules, which could enhance the functional diversity of these regions and provide a network foundation for integrative information processing.

The interdependent connectome-cognitive association analysis results validated our hypothesis that the transmodal cortex plays a coordinating role between the interactive SC and FC networks, thereby promoting cognitive diversity. We found that higher-order cognitive functions were significantly correlated with multilayer modular variability in the transmodal cortex but not in the low-order cortex. This raised the possibility that the SC-FC correspondence in the high-order cortex provides a network-level basis for meeting the high cognitive demands of the human brain. Our result was consistent with previous findings that the transmodal cortex possesses circuit properties essential for human cognition and supports high-level cognitive processes ^34, 62, 63^.

Interestingly, negative loadings were observed in some transmodal regions, most notably the medial prefrontal cortex and posterior cingulate cortex. This implied that lower multilayer modular variability in these regions is associated with better cognitive performance. Recent research has yielded similar results: the medial prefrontal cortex and posterior cingulate cortex showed strong SC-FC coupling, and SC-FC coupling of the posterior cingulate cortex is associated with executive function performance ^4^. A plausible hypothesis arising from these observations is that the tight coupling of the SC and FC enables these regions to maintain a relatively consistent module configuration in SC and FC, thereby supporting cognitive demands. The tight module correspondence in these regions provides efficient communication for other areas that are highly interconnected within the transmodal cortex, thereby promoting better cognitive performance.

The similarity of the overall multilayer modular topography between MZ pairs was greater than that between DZ and sibling pairs, suggesting that multilayer modular variability is under genetic control. Our analysis revealed substantial regional heterogeneity in the heritability of multilayer modular organization. The lateral frontal and inferior parietal regions with higher multilayer modular variability demonstrated relatively relaxed genetic control. As important components of the distributed association cortex, the parietal and frontal cortices undergo protracted maturation processes during human development and are therefore exposed to environmental factors more than sensory regions ^34^. The relatively low heritability of multilayer modular variability observed in these regions could be due to their greater sensitivity to environmental influences. Taken together, our findings demonstrated the extent to which the module relationship between the interactive SC and FC connectome is influenced by genetics.

Recent studies have shown that neuromodulatory systems play an essential role in understanding how the fixed human anatomical connectome can give rise to rich brain functions, in which neuromodulatory systems can dynamically modulate the brain connectome to enable rich behaviors ^35, 64-67^. Neurotransmitters are important components of the brain’s molecular organization and extensively influence synaptic transmission within neural circuits ^68^. Previous studies have shown that receptor distributions reflect the organization of brain connectomes ^36^ and that neurotransmitters coordinate dynamic interactions between modules ^35^. Here, we further demonstrated that the multilayered modular organization of the interdependent structural-functional connectome is also associated with the distributions of multiple receptors, such as MOR, CB_1_, 5-HT_4_, *α*_*4*_*β*_*2*_ and 5-HT_1B_, and is modulated by multiple neurotransmitters, such as opioid, cannabinoid, serotonin and acetylcholine. Previous work has shown that neurotransmitter receptor density distribution forms a natural axis in the human cerebral cortex, extending from sensory to association areas, with association areas having more receptor expression and greater synapse density ^36, 69^. This may provide the anatomical basis for neurons in areas of high multilayer modular variability to integrate information. These neurotransmitters associated with multilayer modular variability are also thought to support many cognitive functions.

Acetylcholine is often implicated in attention control ^70-72^. Enhancing or impairing cholinergic activity can preferentially affect the maintenance of selective attention ^73^. Serotonin (5-hydroxytryptamine, 5-HT) is widely distributed throughout the brain and is involved mainly in learning and memory processes ^74^. Among its receptor families, the 5-HT_1B_ receptor is located predominantly at axon terminals and facilitates learning when cognitive demands are high. In addition, acetylcholine and serotonin are essential for maintaining synapses in the hippocampus and thus play important roles in the acquisition of spatial memory ^75, 76^. Opioids ^77^ and cannabinoids ^78^ are also involved in a wide range of cognitive activities. Taken together, these findings revealed a prominent link between receptor distribution and the multilayer modular architecture of the interdependent structural-functional connectome.

In addition to receptor density, gene expression provides critical neurobiological insight into the function and structure of the brain ^30, 79^. Building on previous reports linking gene expression to modular architecture ^32^, we mapped gene expression patterns to the multilayer module organization in the interdependent structural-functional connectome. We identified a significant association between gene transcription and multilayer module topography and found that genes associated with multilayer module variability are mainly responsible for the biological processes of chemical synaptic transmission. Synaptic transmission is important for supporting the propagation of signals between neurons, and this process is highly dependent on neurotransmitter systems. The presynaptic neuron releases neurotransmitters into the extracellular space via exocytosis of vesicles, and these neurotransmitter molecules are subsequently transported through chemical synapses and bind to appropriate receptors postsynapse ^80^. Thus, this process modifies the neural states of postsynaptic neurons and ultimately results in network-wide communication. Our study showed that genes involved in signal propagation have higher expression levels mainly in regions with higher multilayer modular variability, which has implications for the higher communication demands in these regions.

Several methodological issues need to be mentioned. First, we used a common type of multilayer network in which the SC and FC were connected only via interlayer edges between a given node and its counterparts in other layers to reflect internetwork interactions. The cross-modality couplings between different brain regions were not considered, as there is no generally accepted approach for such an analysis. Therefore, future studies should develop a new strategy to characterize multilayer networks consisting of more complex interlayer connections. Second, we used dMRI-based tractography algorithms to generate representations of white matter tracts in the human brain. However, there are inherent limitations in inferring a reliable SC from fiber tractography approaches ^81, 82^, such as the potential to underestimate long-range connections in the whole-brain network ^83^ and the possibility of missing some short fiber bundles ^81^. Therefore, methodological innovations to reduce tractography biases and improve the reliability of SC-FC estimation are needed in future studies. Third, the gene expression maps used in our study were derived from the Allen Institute for Brain Science; therefore, our current findings are based on a small sample of postmortem brains. In the future, the availability of more comprehensive microarray gene expression datasets will be essential. Finally, our study focused on multilayered modular reconfiguration across the SC and FC layers in healthy participants. Future studies could further investigate whether and how the interactive structural-functional connectome changes with disease, in particular identifying the nodes responsible for communication between these two networks and whether these nodes undergo role changes in patients with brain disorders. It would also be interesting to investigate the age-related changes in the interdependent relationship between the SC and FC.

## Methods

### Participants and data acquisition

The multimodal neuroimaging data (structural MRI, dMRI, and rs-fMRI data) were obtained from the publicly available S1200 dataset released by the HCP ^37^. The HCP S1200 dataset included 1,012 healthy young adult participants (aged 28.73 ± 3.71 years, 543 females) with complete minimally preprocessed imaging data for all modalities. For each participant, there were four rs-fMRI scans (the data were collected over two days; individuals were scanned twice a day (left-to-right and right-to-left phase encoding directions)) and one complete dMRI scan. All functional and diffusion imaging data were preprocessed using HCP minimal preprocessing pipelines ^84^. The HCP obtained informed consent from all participants. The scanning protocol was approved by the Institutional Review Board of Washington University in St. Louis, MO, USA (IRB #20120436).

Structural, functional, and diffusion MRI data were acquired on a 3T Siemens Skyra scanner at Washington University. Specifically, for each run of four rs-fMRI scans for each participant, the rs-fMRI data were obtained by using multiband gradient-echo-planar imaging with the following sequence parameters: repetition time (TR) = 720 ms, echo time (TE) = 33.1 ms, flip angle = 52°, bandwidth = 2290 Hz/pixel, field of view = 208 × 180 mm^2^, matrix = 104 × 90; 72 slices, voxel size = 2 × 2 × 2 mm^3^, multiband factor = 8, and 1200 volumes. Diffusion data from each participant were acquired by using a Stejskal-Tanner diffusion-encoding scheme with the following sequence parameters: 1.25 mm isotropic, 18 b0 acquisitions, 270 diffusion-encoding directions with three shells of b = 1000, 2000, and 3000 s/mm^2^, 90 directions for each shell, 2 × 2 × 2 mm isotropic voxels, TR = 5520 ms, and TE = 9.58 ms. T1-weighted image data were acquired using a 3D-magnetization-prepared rapid acquisition with gradient echo (MPRAGE) sequence (0.7 mm isotropic voxels, matrix = 320 × 320; TR = 2400 ms, TE = 2.14 ms, 256 slices, and flip angle = 8°). T2-weighted data were acquired using a 3D T2-sampling perfection with application-optimized contrasts using a flip angle evolution (SPACE) sequence with identical geometry (TR = 3200 ms and TE = 565 ms).

### Data preprocessing

T1-weighted and T2-weighted images were processed using the minimal structural preprocessing pipeline ^84^, which included brain tissue segmentation, cortical surface reconstruction, and individual surface mapping to the fs_LR_32K standard space. All functional MRI data, including gradient distortion correction, head motion correction, echo-planar imaging distortion correction, registration to the Montreal Neurological Institute (MNI) space, and intensity normalization, were preprocessed. The volume time series of cortical gray matter were then projected onto the standard 32K_fs_LR mesh. A 2-mm full-width at half-maximum (FWHM) Gaussian kernel was used for spatial smoothing. The ICA-FIX procedure was used to remove additional noise. The confounding covariates white matter, cerebrospinal fluid, global signals, and the 12 head motion parameters were further regressed from the time course of each voxel. Finally, bandpass filtering (0.01–0.1 Hz) was performed to reduce the influence of low-frequency drifts and high-frequency physiological noise. The above procedures were carried out using SPM12 (https://www.fil.ion.ucl.ac.uk/spm/) and GRETNA ^85^. The diffusion images were normalized to the mean b0 image, with echo planar imaging (EPI) distortion correction, eddy-current distortion correction, head motion correction, gradient nonlinearity distortion correction, linear registration to native structural space using a 6 degrees of freedom (DOF) boundary-based registration, and data masking with the final brain mask to reduce the file size.

### Constructing interdependent structural and functional connectome

#### (i) Functional connectome (FC)

Based on the preprocessed rs-fMRI data, we constructed the FC of each run for each participant. Specifically, we used a multimodal brain atlas (HCP-MMP1.0) to parcellate the cortical surface into 360 areas ^42^. The time series of all vertices within each node were averaged to generate the mean time series of each node. Pairwise Pearson correlations were then calculated between the mean time series of all nodes to generate functional connectivity edges. As a result, we obtained a Pearson correlation matrix of size 360 × 360 for each run for each participant. To reduce signal noise bias, the weak connections (Pearson’s r < 0.1) of each correlation matrix were set to zero. We also validated different weak connection thresholds of the FC matrices (Supplementary Figs. 6 and 7). Finally, Fisher’s r-to-z transformation was applied to each FC matrix. For each participant, the FC matrices of all four rs-fMRI scans were averaged to generate the mean FC matrix.

#### (ii) Structural connectome (SC)

Structural connectivity was estimated for participants using probabilistic tractography. The analysis procedures were implemented in the FSL ^86^ and the PANDA Toolkit ^87^. Specifically, for a given seed region, probabilistic tractography was performed by sampling 5,000 streamline fibers for each voxel within that region. According to the number of streamlines between the source and target regions, the connectivity probability between these two regions was calculated as the number of streamlines passing through the target region divided by the total number of streamlines sampled from the seed region. Notably, the long-range connections may be underestimated due to the fact that the number of streamlines decreases with distance from the seed mask. Therefore, distance correction was then applied to obtain connectivity weights between regions; these weights were defined as the expected length of the paths times the number of streamlines ^87, 88^. Using the above procedure, we obtained the connectivity weights for all pairs of brain nodes, resulting in an SC matrix of size 360 × 360 for each participant.

#### (iii) Interdependent structural□*functional connectome*

Using the multilayer network theory ^11, 12, 89^, we modeled the interdependencies between the SC and FC. Specifically, the SC and FC connectomes were considered separate layers of an interdependent network. The different layers shared the same set of nodes, with the number of nodes in each layer equal to 360. We then established internetwork dependencies by multiplex coupling, in which the corresponding nodes located in different layers were coupled in a one-to-one manner, generating a two-layer interdependent structural-functional connectome for each participant, which can be represented by a supra-adjacency matrix ^44^ ℳ with the following form:

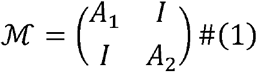

where *A*_*i*_ is the adjacency matrix of layer 𝓁_*i*_(*i* = 1,2). *I* is the *N* × *N* (N = 360) identity matrix. Inherent discrepancies in weight scales between different modalities can lead to biases in multilayer network analysis ^90^. In the present study, the SC matrix had significantly larger weights than did the FC matrix. This discrepancy posed the risk that the SC layer would disproportionately influence the multilayer modularity detection algorithm, potentially biasing the resulting modules to predominantly reflect SC features. To ensure balanced contributions from each layer, we normalized the weights of both the SC and FC matrices to a uniform range of 0-1.

### Identifying multilayer connectome modules

Modularity is an important organization principle for brain connectomes ^24, 26, 91^. The existence of modules allows the brain to achieve effective information communication at low wiring costs ^92^. In the context of a multilayer network, we used a generalized Louvain-like locally greedy algorithm (https://github.com/GenLouvain/GenLouvain) to obtain the multilayer modular architecture by simultaneously considering all the information within and between layers ^38, 39^. The main idea of this GenLouvain community detection algorithm is to optimize the multilayer modularity quality function Q to identify the module membership of each node in the network. The modularity quality function of the interdependent structural-functional connectome was calculated as follows:

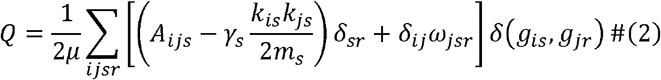

where *μ* represents the total connectivity strength of the entire network. Nodes are represented by i and j. Layers are represented by s and r. *A*_*ijs*_ is the element of the correlation matrix and represents the connectivity strength between node i and node j in layer s. *k*_*is*_ and *k*_*js*_ are the degrees of node i and node j in layer s, respectively. *m*_*s*_ represents the total connectivity strength of layer s. The result of *k*_*is*_*k*_*js*_/2*m*_*s*_ reflects the expected connection probability between node I and node j in layer s. *g*_*is*_ indicates the module in which node i belongs in layer s. *g*_*j*_ indicates the module in which node j belongs in layer r. The function *δ*(*g*_*is*_, *g*_*jr*_) is used to determine whether the modules of node i and node j are the same. *δ*(*x, y*) equals 1 if x = y and equals 0 otherwise. The interlayer coupling parameter *ω* reflects interlayer dependence. A higher *ω* indicates a stronger interaction between layers, and vice versa. The topological resolution of each layer is represented by the parameter *γ*. A larger value of *γ* indicates a larger number of modules. Since there is currently no uniform standard for the choice of parameters *γ* and *ω*, we used the default value of *γ* = *ω* = 1, as was done in previous studies^93-95^. We also calculated multilayer modularity using different interlayer coupling parameters with *ω* (*ω* = [0.5, 0.75, 1, 1.25, 1.5]) (Supplementary Fig. 8).

By optimizing the modularity quality function (2) with a Louvain-like locally greedy heuristic algorithm ^39^, we obtained the multilayer modular structure of the interdependent structural-functional connectome. However, since this algorithm is heuristic in nature and due to the near degeneracy of the optimization landscape of the multilayer modularity quality function, the results of multilayer modularity detection may be slightly different each time the algorithm is run ^62, 96, 97^. Therefore, to address the issue of degeneracy, we repeated the Louvain-like locally greedy algorithm 100 times for each participant to optimize the modularity index Q. Based on the results of each run, we calculated the corresponding multilayer network measurements and finally averaged the results of the 100 runs for each participant.

### Tracking multilayer modular variability

The multilayer modularity detection algorithm ^38, 39^ was used to extract the modular architecture of the interdependent structural-functional connectome, generating module assignments for both the SC and FC layers. To investigate the correspondence of module organization between interdependent SC and FC layers, we tracked the modules of each node across layers. Each node was identified with community labels, which may be consistent or inconsistent across different layers. Therefore, we evaluated the cross-layer module affiliation variability of nodes using the multilayer modular variability (MV) metric ^40^. For a given node i in the network, the multilayer modular variability of that node was calculated as follows:

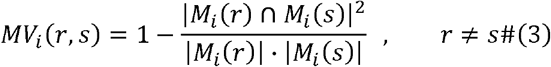

where r and s represent layer labels and *M*_*i*_(*r*) *M*_*i*_(*s*) abels of the modules in which node i belongs in layers r and s, respectively. |*M*_*i*_(*r*)| denotes the number of nodes included in module *M*_*i*_(*r*). |*M*_*i*_(*r*) ⋂ *M*_*i*_(*s*)| indicates the number of overlapping nodes between modules M_*i*_ crJ and M_*i*_ csJ. The multilayer modular variability reflects the degree of spatial variability in the module organization across the SC and FC layers. A node with high (low) multilayer modular variability has a large (small) difference in module organization across the SC and FC layers.

### Reliability and reproducibility analysis of multilayer modular variability

#### (i) Reliability analysis

We used the HCP Test-Retest dataset to quantitatively assess whether the multilayered modular variability pattern in the interdependent structural-functional connectome was reliable within participants across repeated sessions and variable between participants. The HCP Test-Retest dataset included 42 participants (aged 30.4 ± 3.33 years, 30 females) who underwent a second MRI scan (second session: S2) between 0.5 and 11 months after the first scan (first session: S1). The data acquisition, data preprocessing, and connectome construction methods used for the HCP Test-Retest dataset were consistent with those used for the HCP S1200 dataset. For a given participant, we separately constructed the interdependent structural-functional connectome and calculated the multilayer modular variability of that participant for two sessions (i.e., S1 and S2). We then calculated the Pearson correlation coefficient for each participant’s multilayer modular variability obtained between S1 and S2, which was considered an indicator of within-participant similarity. In addition, we assessed between-participant similarity by calculating the mean Pearson correlation between each participant’s multilayer modular variability in S1 and that of all the other participants in S2. To determine whether there was a significant difference between these within-participant and between-participant similarities, we performed a nonparametric permutation test, randomizing participant identities over 10,000 repetitions. We recalculated the similarities and generated null distributions for the differences in similarity. The statistical significance of the observed difference in similarity was then assessed by comparison with this null model.

Furthermore, we used the intraclass correlation (ICC) ^49^ to examine the test-retest reliability of the multilayer modular variability in each brain region across repeated sessions. Specifically, for a given brain region, the multilayer modular variability of that region can be combined into a matrix M, where the element M_ij_ represents the multilayer modular variability from jth measurement session of the i-th participant. A one-way analysis of variance was then performed on this matrix to obtain the within-participant mean square error (*MS*_*W*_) and the between-participant mean square error (*MS*_*b*_). The ICC of the given region can be calculated as follows:

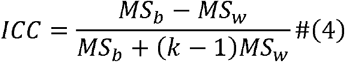

where k is the number of repeated sessions for each participant. High ICC values reflect low within-participant variance relative to between-participant variance. According to previous study ^98^, the ICC values can be divided into four common intervals: poor (< 0.4), moderate (0.4∼0.6), good (0.6∼0.75), and excellent (> 0.75).

#### (ii) Reproducibility analysis

To investigate whether the multilayer modular topography in the interdependent structural-functional connectome is reproducible, we employed a half-split strategy to randomly (n = 1,000 repetitions) divide the 1,012 participants into two subgroups. For each random half-split result, we used the chi-square test and two-sample t test to ensure that the two subgroups were matched for sex and age, respectively. Next, we calculated the group-level multilayer modular variability pattern of each subgroup and further evaluated the Pearson correlation between these patterns to estimate the reproducibility of the multilayer modular variability. Fig. 3c shows the results of one of the 1,000 random divisions (subgroup 1: 506 participants, aged 28.70 ± 3.66 years, 272 females; subgroup 2: 506 participants, aged 28.76 ± 3.77 years, 271 females).

### Heritability analysis

To investigate whether the multilayer modular topography in the interdependent structural-functional connectome is influenced by genetic factors, we conducted a similarity analysis of multilayer modular variability. Based on the twin and family data in the HCP S1200 dataset (n = 1,012 participants), we determined the zygosity of the participants using genotyping data when available and self-reports otherwise. Three participants were excluded due to abnormal family data. The final sample consisted of 1,009 participants from 449 families, including 268 MZ twins, 140 DZ twins, 107 singletons, and 494 nontwins. We compared whether the interindividual similarity in multilayer modular variability differed among MZ twins, DZ twins, and nontwins. Briefly, we calculated the similarity (Pearson’s r) in overall multilayer modular variability between pairs of participants, assessing the extent to which the multilayer modular organization of two participants became more similar as their proportion of shared genetic material increased, where r_MZ_ was the correlation between MZ twins, r_DZ_ was the correlation between DZ twins, and r_nontwin_ was the correlation between nontwins. We then used a nonparametric permutation test (10,000 repetitions) to estimate whether there were differences in the r_MZ_, r_DZ_ and r_nontwin_ data. For each permutation test, participant identities were randomly shuffled, and the r_MZ_, r_DZ_ and r_nontwin_ values were recomputed, generating the null distribution of the mean difference of r_MZ_-r_DZ_, r_DZ_-r_nontwin_ and r_MZ_-r_nontwin_. The statistical significance of the mean difference was calculated by comparing the observed value with the null model.

To further investigate the extent to which genetic factors underlie the spatial layout of multilayer modular organization in the interacting structural-functional connectome, we applied the accelerated permutation inference for the ACE model (APACE) method ^99^ (https://github.com/nicholst/APACE) to estimate the heritability of multilayer modular variability. The ACE model of heritability analysis relies on the assumption that phenotypic variability within a population can be explained by additive genetic (A), common environmental (C) and unique environmental (E) factors. The APACE model mainly relies on linear regression with squared differences to estimate phenotypic heritability and the likelihood ratio test to infer heritability. Heritability represents the proportion of phenotypic variation attributable to genetic variation ^100^. Narrow-sense heritability (*h*^2^) was calculated as follows:

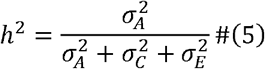

where 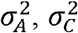 and 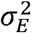 are the variance of A, C and E, respectively.

### Multilayer connectome-cognition association analysis using partial least squares regression

We used PLS regression analysis (https://github.com/danizoeller/myPLS) to explore how the spatial distribution of multilayer modular variability in the interdependent structural-functional connectome corresponds to cognitive processes. Using all the cognitive data provided by the HCP S1200 dataset, we performed PLS analysis to decompose the relationships between multilayer modular variability (dataset X: 1,012 participants × n brain regions) and cognition (dataset Y: 1,012 participants × 52 cognition terms) into orthogonal sets of latent variables with maximum covariance ^101^. These latent variables were linear combinations of the original data from the two datasets and consisted of singular vectors and singular values. Specifically, datasets X and Y were z scored column by column, and the covariance matrix R was subsequently calculated:

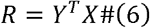

Next, singular value decomposition (SVD) was performed on R:

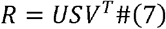

where U and V are the left and right singular vectors, respectively, and S is a diagonal matrix of singular values. The ith latent variable is composed of the ith left and right singular vectors and the ith singular value. The ith singular value represents the covariance between X and Y that is captured by the corresponding ith latent variable. According to the singular value, we estimated the amount of covariance explained by each latent variable, which is the ratio of the square of the ith singular value to the sum of the squares of all singular values. The left and right singular vectors represent the cognitive weights and multilayer modular variability weights, respectively, which reflect the extent to which each cognitive term and brain region contribute to the latent variable. By projecting the original data onto the weights of the singular vectors, we obtained the cognitive scores and multilayer modular variability scores. Pearson correlation analysis between cognitive scores and multilayer modular variability scores was conducted to characterize the relationship between multilayer modular variability and cognition under the given latent variable. Furthermore, we correlated each original variable with its PLS analysis-derived score pattern to compute the cognitive loadings and multilayer modular variability loadings, which reflect the shared variance between the original variables and their corresponding score pattern and further reveal the degree of contribution of the cognition terms and brain regions to the corresponding latent variable.

To assess the statistical significance of each PLS latent variable, we performed a nonparametric permutation test by randomly shuffling the rows (participant identities) of matrix Y 10,000 times. For each permutation test, we recalculated the covariance matrix R and performed SVD. Consequently, we obtained the null distribution of the singular values and the PLS score patterns. By comparing the empirical values with their null distributions, we estimated the statistical significance of each latent variable. In addition, the reliability of the loadings of the variable was assessed using bootstrapping resampling (1,000 repetitions). We conducted a bootstrapping analysis by randomly resampling participants. Using the resampled data matrices X and Y, we performed SVD again and recalculated the loadings of the variables. According to the 95% CIs of the variable loadings, we selected the brain regions and cognitive terms that made significant contributions to the latent variable.

### Multilayer connectome-transmitter association analysis using elastic net regression

Building on previous work showing that brain modules are closely associated with neurotransmitter systems ^35, 36^, we investigated whether the multilayered module organization in the interdependent structural-functional connectome is also supported by the underlying molecular mechanisms involved. Following a previous study ^36^ that provided cortical distribution data of 19 neurotransmitter receptors and transporters from nine neurotransmitter systems, we calculated the average density of each cortical region in the Glasser360 atlas for each of the 19 receptors and transporters, resulting in a density matrix of size 360 × 19. We then used a multivariate elastic net regression model to predict multilayer modular variability from the receptor and transporter density distributions. As it is a data-driven regression approach, multivariate elastic net regression can be used to solve the multicollinearity problem between independent variables and can be used for feature selection by automatically removing variables that are deemed unrelated to the dependent variable, resulting in a sparse output. Therefore, this method was well suited for receptor and transporter data where variables are highly correlated with each other (Supplementary Fig. 9), and some variables may be less important for fitting multilayer modular variability. As the numerical scales of the receptor and transporter data and the multilayer modular variability data differed, all variables were normalized prior to regression analysis. For a given variable s (360 brain regions × 1), the normalization process was as follows:

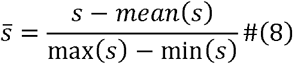

Furthermore, multivariate elastic net regression was used to fit multilayer modular variability. This approach is based on the loss function of the model by adding a penalty term consisting of L1 regularization (LASSO regression) ^102^ and L2 regularization (Ridge regression) ^103^. L1 regularization is used to penalize the sum of the absolute values of the model parameters, excluding features with smaller contributions to the target variable, thus achieving feature selection. L2 regularization is used to penalize the sum of squares of the model parameters to achieve weight decay (i.e., nonsparse *β* values), which helps to prevent multicollinearity problems and reduce the model’s overreliance on certain features. The objective function of the multivariate elastic net regression ^104^ is as follows:

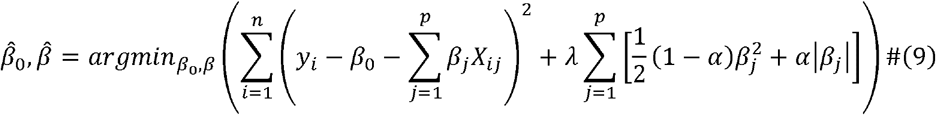

where n and p represent the number of samples (n = 360) and features (p = 19), respectively. X and y, respectively, represent the normalized receptor and transporter data and the normalized multilayer modular variability data. *β*_0_ is the intercept, and *β*_*j*_ is the regression coefficient of the jth feature. *α* represents the mixed ratio of L1 (*α* = 1) and L2 (*α* = 0) regularization. To combine the advantages of Ridge regression, which can address multicollinearity, and LASSO regression, which can be used to perform feature selection, we set the *α* value to 0.5 ^104^. The regularization coefficient *λ* is used to control the intensity of the penalty and to determine the sparsity of the model output. An optimal *λ* value can be selected through a cross-validation model. Specifically, we divided the range from 10^−5^ to 10^2^ into 100 equal parts, and these 100 values were considered a selectable range of parameters, denoted as *λ*. For each *λ* value, multivariate elastic net regression analysis was conducted, and 10-kold cross-validation was used to evaluate model performance. Finally, the *λ* value with the lowest mean square error was selected as the optimal parameter.

To test whether the real R^2^ of the model was significantly greater than that obtained by chance, we performed a spin test 10,000 times, generating 10,000 null distribution maps of multilayer modular variability. For each surrogate multilayer modular variability map, we conducted multivariate elastic net regression governed by the optimal regularization parameter *λ* from the empirical model to predict multilayer modular variability, resulting in a null distribution of model R^2^ to test the statistical significance of the empirically observed model R^2^.

### Multilayer connectome-transcriptome association analysis using partial least-squares regression

#### (i) AHBA gene expression dataset

To investigate the associations between spatial configurations of the multilayer module structure in the interdependent structural-functional connectome and transcriptional profiles, we used the microarray data of six human postmortem donors (aged 42.5 ± 13.38 years, 1 female) provided by the AHBA website (http://human.brain-map.org/) ^41^ to estimate gene expression in the brain. A total of 3,702 spatially distinct tissue samples were obtained from the six donors, and 58,692 probes were obtained for each sample. Since sample data from the right hemisphere were available for only two donors, we analyzed the tissue samples from the left hemisphere only.

#### (ii) Preprocessing of gene expression data

Following the AHBA processing pipeline (https://github.com/BMHLab/AHBAprocessing) ^105^, we preprocessed the microarray-based gene expression data collected from human brain tissue samples from six adult donors. Specifically, the probe-to-gene annotations were updated using the data provided by Arnatkeviciute et al ^105^. We then removed the probes whose signal-to-noise ratio did not exceed the background noise by using intensity-based filtering with a threshold of 0.5. Considering the difference between probes that measure the expression of the same gene, we selected the probes that exhibited the strongest correlation with the RNA-seq data. Next, using the MNI coordinates of each tissue sample, we assigned each sample to the region nearest to the Glasser360 brain parcellation with a distance threshold of 2 mm. Tissue samples more than 2 mm away from any region of the 360-parcellation were excluded. To account for interindividual variability in gene expression, we applied the scaled robust sigmoid normalization method to the left cortex data to eliminate this donor-specific variability. The normalization procedures were first performed by applying cross-gene normalization in a given sample. Then, cross-sample normalization was performed for each gene. Finally, all samples from six donors were averaged for a given region, resulting in a group-level gene expression matrix of size n (132 brain regions) × g (10,027 genes).

#### (iii) Association between multilayer modular variability and transcriptional signatures

We assessed the relationship between multilayer modular variability and gene expression using multivariate PLS regression. The gene expression matrix (132 brain regions × 10,027 genes) and multilayer modular variability (132 brain regions × 1) were considered predictor variables and response variables, respectively. The PLS regression method was used to attempt to find the PLS components that are linear combinations of the original gene expression that can maximize the prediction of the response variables. We calculated the R^2^ of the model fitting, which reflects the amount of variance in multilayer modular variability explained by each PLS component. In addition, the Pearson correlation was conducted to estimate the spatial correlation between the PLS scores and the multilayer modular variability map. To assess whether the empirical R^2^ and Pearson’s r values were significantly greater than those obtained by chance, spatial autocorrelation correction (spin test) was performed, generating 10,000 null distribution maps of the multilayer modular variability. For each permutation, the real predictor variables and the surrogate response variable were assessed by PLS regression analysis, and we recalculated the R^2^ of each PLS component, generating a null distribution of variance explained. Similarly, a null distribution of correlation coefficients (r) between the PLS score and multilayer modular variability under each PLS component can be obtained. The P value (i.e., P_spin_) was calculated as the proportion by which the values (i.e., R^2^ or Pearson’s r) of the null models were greater than the empirically observed values.

#### (iv) GO enrichment analysis

To explore the enriched GO terms associated with genes identified by PLS analysis, we performed GO enrichment analysis by using the online tool GOrilla (http://cbl-gorilla.cs.technion.ac.il/) ^106^. First, for each significant PLS component, we calculated the contribution weights of the genes and assessed the reliability of the weights by bootstrapping resampling (1,000 repetitions). For each resampling, the rows of the gene expression matrix were randomly selected to generate the new bootstrapped gene expression matrix, which was used when PLS analysis was performed again. This process was repeated 1,000 times to obtain a sampling distribution of gene weights, and we further estimated the standard errors of these weights. We then computed the bootstrap ratio ^107^ of the genes by dividing the empirical weights by their standard errors, with large bootstrap ratios representing the genes with large and reliable contributions. Thus, we generated a gene list for each PLS component to represent the contribution of the genes. Furthermore, we ranked the gene list in both descending and ascending order and subjected these ranked gene lists to the GOrilla software tool to search for enriched GO terms for each PLS component. The significant enrichment terms were identified by applying the FDR-corrected q < 0.05. With respect to the advanced parameter settings of the GOrilla platform, we selected the “P value threshold 10^−4^” and unchecked the “Run GOrilla in fast mode” option ^108, 109^. Finally, we used the Reduce Visualize Gene Ontology (REVIGO, http://revigo.irb.hr/) tool to summarize these significant GO terms by removing redundant GO terms.

### Spatial autocorrelation-preserving permutation tests

The spatial autocorrelation-preserving permutation test, which applies random rotations to spherical representations of the cortical surface, is also known as the spin test ^110^. Briefly, we mapped the spatial distribution of multilayer modular variability in the interdependent structural-functional connectome onto the cortical surface, and multilayer modular variability value was obtained for each vertex. The spin test was applied to generate 10,000 rotational permutations of multilayer modular variability. For a given node, the surrogate multilayer modular variability value was assigned as the mean value of the vertexes within that node. As a result, surrogate brain maps of multilayer modular variability were generated to assess the statistical significance of the spatial correspondence between multilayer modular variability and cortical gradient, cortical expansion, ICC, neurocognitive flexibility, neurotransmitter receptor and transporter density distributions, and gene expression.

In addition, the spin test was used to assess whether the mean multilayer modular variability, ICC and heritability of each hierarchical system were determined by the cortical partitions or by spatial autocorrelation. Briefly, we performed the spin test (10,000 repetitions) to permute the positions of the systems under the premise of preserving spatial autocorrelation, and then the mean multilayer modular variability, ICC and heritability value for each system were recomputed. These mean values were further expressed as z scores relative to the null model, with positive/negative z values representing real values greater/smaller than those expected by chance. The P value (i.e., P_spin_) of the spin test was calculated as the proportion by which the values of the null models are greater in magnitude than the empirical observations.

## Reporting summary

Further information on research design is available in the Nature Research Reporting Summary linked to this article.

## Data availability

The HCP dataset, including structural MRI, functional MRI, and diffusion-weighted MRI, is available in the HCP ConnectomeDB (https://db.humanconnectome.org/). The neurocognitive flexibility data is publicly available at https://surfer.nmr.mgh.harvard.edu/fswiki/BrainmapOntology_Yeo2015. The neurotransmitter receptor and transport density distribution data are publicly available at https://github.com/netneurolab/hansen_receptors. The AHBA dataset is publicly available at https://human.brain-map.org/static/download. Intermediate data supporting the results are available at https://github.com/wangxyue/Topographic-cognitive-neurobiological-profiling-of-interdependent-SC-FC.

## Code availability

All analysis code is available at https://github.com/wangxyue/Topographic-cognitive-neurobiological-profiling-of-interdependent-SC-FC.

## Acknowledgements

This study was supported by the National Natural Science Foundation of China (Nos. 82021004) and Beijing Natural Science Foundation (JQ23033). We thank Dr. Tianyuan Lei for the discussion on the experimental design. The imaging data were provided by the Human Connectome Project, WU-Minn Consortium (Principal Investigators: David Van Essen and Kamil Ugurbil; 1U54MH091657) funded by the 16 NIH Institutes and Centers which support the NIH Blueprint for Neuroscience Research; and by the Mc-Donnell Center for Systems Neuroscience at Washington University.

## Author contributions

Xiaoyue Wang and Lianglong Sun performed all analyses and wrote the manuscript. Xinyuan Liang helped with the data processing and analysis of gene. Mingrui Xia contributed to the experimental design and interpretation of experimental results of the genetic association analysis. Tengda Zhao assisted with data processing and analysis of neurotransmitter receptors and transporters. Xuhong Liao provided assistance in experimental design, analysis of multilayer network and interpretation of experimental results. Yong He designed and supervised the study, developed the main concepts, interpreted results, contributed to the writing, reviewing and editing of the manuscript and secured funding.

